# Adaptive mechanisms facilitate robust performance in noise and in reverberation in an auditory categorization model

**DOI:** 10.1101/2022.09.25.509412

**Authors:** Satyabrata Parida, Shi Tong Liu, Srivatsun Sadagopan

## Abstract

For robust vocalization perception, the auditory system must generalize over variability in vocalization production as well as variability arising from the listening environment (e.g., noise and reverberation). We previously demonstrated that a hierarchical model generalized over production variability by detecting sparse intermediate-complexity features that are maximally informative about vocalization category from a dense spectrotemporal input representation. Here, we explore three biologically feasible model extensions to generalize over environmental variability: (1) training in degraded conditions, (2) adaptation to sound statistics in the spectrotemporal stage and (3) sensitivity adjustment at the feature detection stage. All mechanisms improved vocalization categorization performance, but improvement trends varied across degradation type and vocalization type. One or both adaptive mechanisms were required for model performance to approach the behavioral performance of guinea pigs on a vocalization categorization task. These results highlight the contributions of adaptive mechanisms at multiple auditory processing stages to achieve robust auditory categorization.

## Introduction

To maintain robust auditory perception, especially of communication sounds such as vocalizations (calls), animals and humans must generalize over the tremendous variability in the production of these sounds and the variability imposed by the listening environment. Production variability includes both trial-to-trial and subject-to-subject variability that is inherent in sound production. Environmental variability refers to a variety of acoustic degradations such as the addition of background noise and reverberation. Call categorization thus requires a many-to-one mapping operation, where diverse acoustic inputs are binned into a small number of behaviorally relevant categories. How this categorization operation is implemented by the neural circuitry of the auditory system is a central question in auditory neuroscience. Existing auditory encoding models are not well-suited to address this question. For example, many auditory pathway models adopt an engineering approach, and process inputs using experimenter-defined spectral, temporal, or modulation filter banks, which are biologically inspired (Chi et al., 2005; Dau et al., 1997); however, these models focus on stimulus encoding and have not been tested in categorization tasks. Another class of models, which are based on deep neural networks, can achieve robust auditory categorization performance, which can approach human performance levels (Kell et al., 2018; Li et al., 2014). However, these complex networks offer limited biological interpretability, and an intuitive understanding of what stimulus features are used to generalize over a given category, and how these stimulus features are biologically computed, is elusive. Additionally, the latter class of models requires large amount of training data (typically of the order of millions of data points). To strike a balance between biological interpretability and categorization performance, we have previously proposed a hierarchical model that learns informative, intermediate-complexity features of call categories that can generalize over within-category variability and accomplish categorization (Liu et al., 2019). This model can achieve robust production invariance with a limited training set (of the order of hundreds of data points). However, in its simplest implementation, model performance degrades when environmental variability is introduced to model inputs. In the present study, we characterize model performance for some common types of environmental degradations and extend the model to include several adaptive neural mechanisms that may aid auditory categorization in such conditions.

The hierarchical model consists of three stages (Fig. 1). The first stage is a dense spectrotemporal representational stage, which is the output of a biophysically realistic model of the cochlear filter bank (Zilany et al., 2014). The second stage consists of a set of sparsely active feature detectors (FDs). Each FD has a spectrotemporal receptive field (STRF), which corresponds to the stimulus feature that the FD is tuned to detect, and a nonlinear threshold. Optimal feature tuning for performing specific call categorization tasks, and optimal thresholds to maximize classification performance, are learned using greedy search optimization and information theoretic principles (Liu et al., 2019). The final ‘voting’ stage of the model obtains the evidence for the presence of a call category by combining the outputs of each category’s FDs, weighted appropriately. Note that, to output the final call category in a go/no-go paradigm, the voting stage can be implemented as a winner-take-all algorithm (Kar et al., 2022). We have previously shown that this hierarchical framework can achieve high accuracy in categorizing calls across multiple species, and that model performance in categorizing natural and manipulated guinea pig (GP) calls mirrors GP behavior (Kar et al., 2022). In addition, we showed that in guinea pigs (GPs), subcortical and layer 4 neurons in the primary auditory cortex (A1) show simple receptive fields and dense spectrotemporal tuning consistent with the spectrotemporal representation stage of the hierarchical model, whereas layer 2/3 neurons in A1 show complex receptive fields and sparse tuning consistent with the FD stage (Montes-Lourido et al., 2021). This observation is consistent with other studies that suggest that receptive field complexity increases between A1 layer 4 and layer 2/3 (Moerel et al., 2019; Montes-Lourido et al., 2021; Sharpee et al., 2011) and that sound category information may be encoded by neurons in the superficial layers of A1 (Bathellier et al., 2012; Xin et al., 2019).

**Figure 1:**
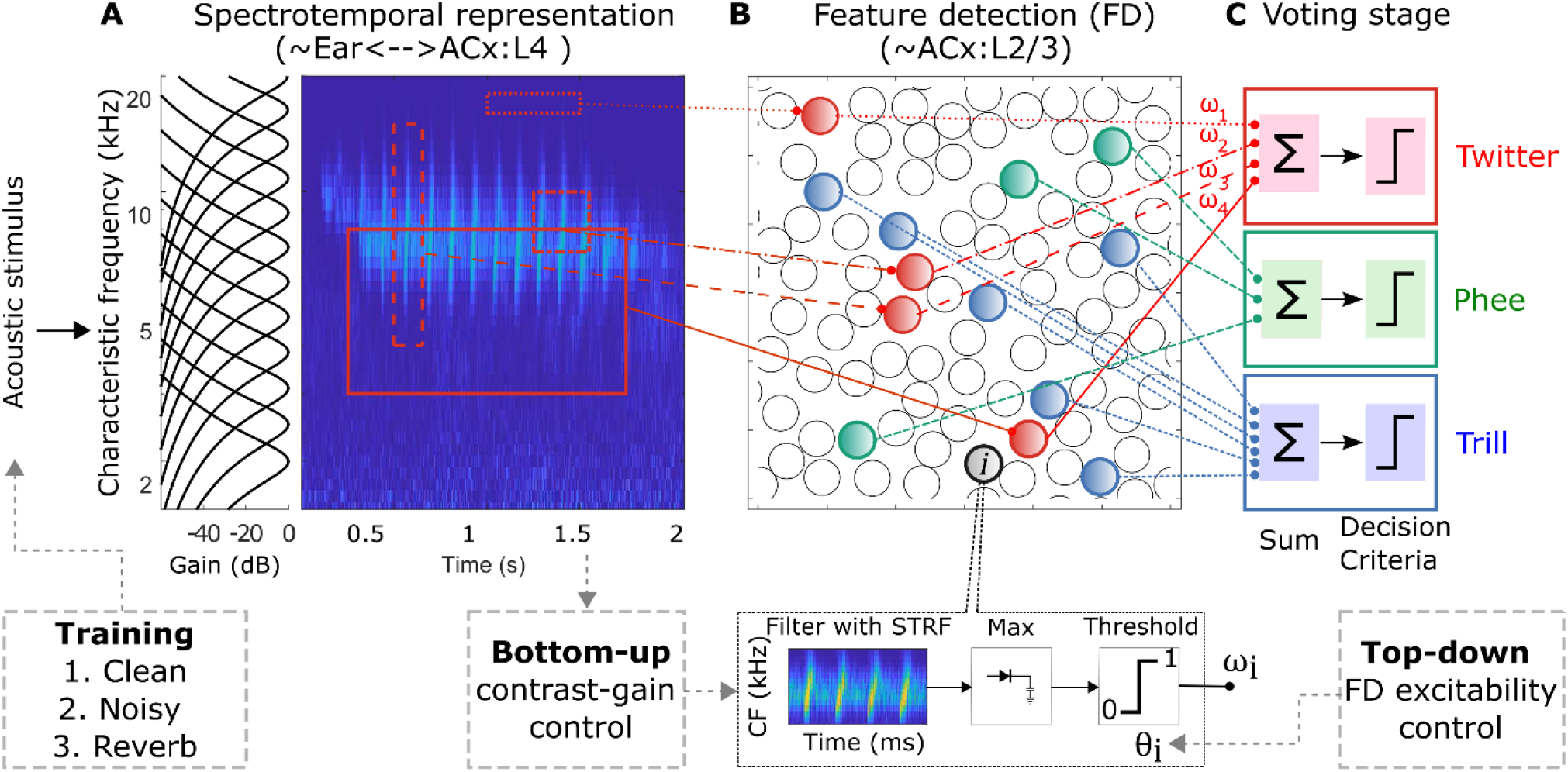
Hierarchical structure of the computational model. The core model consisted of a dense spectrotemporal stage, a sparse feature detection stage, and a voting stage. **(A)** An acoustic stimulus was filtered by a cochlear filter bank to obtain a dense spectrotemporal representation of the stimulus, called a cochleagram. **(B)** The second stage had call-specific feature detectors (FDs) modeled as a spectrotemporal receptive field followed by a threshold. Template matching by cross-correlation (limited to the bandwidth of the FD) was used to obtain a FD *V*_*m*_ response, and the *V*_*m*_ response was compared with the threshold to obtain a binary output. Each call type had a set of maximally informative FDs, whose STRFs and thresholds were learned during model training. **(C)** Finally, in the voting stage, the weighted outputs of the FDs for a given call type were combined to form the final response of the model. In this study, in addition to these stages, we extended the model to include three mechanisms (boxes with dashed lines): **(1)** condition-specific training, **(2)** contrast gain control, and **(3)** top-down modulation of the excitability of FDs. ACx: Auditory cortex; CF, Characteristic frequency; FD, Feature detector; STRF, Spectrotemporal receptive field

But the model is explicitly trained to generalize over the production variability in calls, which can be conceptualized as spectrotemporal variations (limited by biological constraints) around an archetypal call for each category. This variability occurs on a trial-to-trial time scale. In contrast, environmental variability is not a function of call type, is not limited by biological capabilities, and is unlikely to show variations at rapid time scales. Thus, the fundamental strategy employed by the model to generalize over production variability, that of detecting informative features, is unlikely to also confer resilience to environmental variability. Therefore, in this study, we characterized model performance in two challenging conditions: with additive white Gaussian noise and in reverberant settings. We found that the basic model failed to generalize over these forms of environmental variability. Therefore, we extended the model to include several neural mechanisms that are known or hypothesized to improve neural coding in degraded environments. First, condition-specific training can improve the performance of models (Bishop, 1995; Ko et al., 2017) as well as humans and animals (Irvine, 2018). We implemented condition-specific training by using degraded calls (i.e., calls that were affected by noise or reverberation) as the training material. Second, contrast gain control, which refers to adaptive changes to neural tuning and activity levels to match incoming sound statistics, has been demonstrated in several stations along the auditory pathway (Angeloni et al., 2021; Barbour and Wang, 2003; Dean et al., 2005; Lohse et al., 2020; Rabinowitz et al., 2011; Watkins and Barbour, 2008; Wen et al., 2009). We implemented a version of contrast gain control by adapting neural responses in the dense spectrotemporal stage to the mean and standard deviation of population activity in that stage. Finally, recent studies have shown that attention-mediated feedback can control the excitability of cortical neurons to aid performance in challenging listening environments (Fu et al., 2014; Kerlin et al., 2010). Specifically, arousal-related noradrenergic input from locus coeruleus (Maness et al., 2022; Martins and Froemke, 2015) as well as cholinergic inputs from the basal forebrain (Froemke et al., 2007; Kuchibhotla et al., 2017; Letzkus et al., 2011) can increase the excitability of cortical principal neurons, often by providing disinhibition by suppressing the activity of inhibitory interneuron subtypes (Pi et al., 2013). We implemented this top-down pathway as a modulation of the threshold of FDs. While all three mechanisms generally improved model performance both in noise and in reverberation, the trends of benefit varied across degradation type (noise or reverberation) as well as call type, which suggests that the auditory system may differentially rely on these mechanisms based on degradation type and the spectrotemporal properties of the sound. These results demonstrate that multiple adaptive mechanisms acting at multiple auditory processing stages are necessary to achieve robust auditory categorization.

## Results

The computational mechanisms described in this manuscript extend a previously published model of auditory categorization (Liu et al., 2019). For ease of reading, we begin by briefly summarizing some core details of this model (also see Materials and Methods). Model feature detectors were trained to optimally categorize one conspecific call type (within-class) from all other call types (outside-class). To do so, a large number of random features, which served as candidate STRFs that feature detectors (FDs) might detect, were first generated by randomly selecting small portions of within-class cochleagrams with random center frequency, bandwidth, onset time, and duration. For each candidate FD, an incoming cochleagram was convolved with its STRF (restricted to the bandwidth of the STRF) to obtain the membrane potential response (or *V*_*m*_ response). The maximum of the *V*_*m*_ response was compared with the FD’s threshold (learned as described next) to obtain the FD binary response or FD output, where FD output = 1 if maximum of *V*_*m*_ response ≥ threshold. To learn the FD threshold, we first constructed distributions of its *V*_*m*_ response maximum for within-class calls as well as outside-class calls; FD threshold was set to the value that maximized classification merit, as quantified by mutual information between stimulus class and FD output.The FD weight was set to the log-likelihood of classification. From this initial set of ∼5000 candidate FDs, we employed an iterative greedy search algorithm (Ullman et al., 2002) to obtain the maximally informative set of FDs (or MIF set) for the categorization of each call type. To do so, we sequentially added candidate FDs to the MIF set as long as candidate MIFs were not similar (in an information theoretic sense) to existing members of the MIF set and classification performance of the MIF set continued to improve. In the final voting stage, outputs of FDs in the MIF set were weighted (by the learned weight) and summed to obtain the final output of the categorization model. Five separate instantiations of models (with non-overlapping MIF sets) were trained for each call type. Models were tested using a new set of within-class and outside-class call types that were not used in training and model performance was quantified using the sensitivity index, d’.

### Condition-specific training improved model performance but these benefits were typically limited to the same condition

Previous computational and behavioral studies have shown that categorization performance is better when training and testing conditions are the same (e.g., trained and tested in noisy calls) compared to when testing is done in a different condition (e.g., trained with clean calls and tested with noisy calls) (Bishop, 1995; Irvine, 2018; Ko et al., 2017). To determine whether these results also applied to our hierarchical model, we trained and tested the model on calls degraded by noise or reverberation and compared the performance of the noise-trained model to the model trained only on clean calls. To ensure that our results reflected general auditory processing principles, we trained and tested our models on calls from two different species (marmosets and guinea pigs). The calls of these species differ in their fundamental frequencies (marmoset calls have higher fundamental frequencies) and spectrotemporal properties (marmoset calls typically consist of long syllables with continuous frequency contours). Condition-specific training significantly improved model performance both in noise and in reverberation (Fig. 2, Table 1). For example, compared to a model trained on clean calls and tested in noisy calls (black lines in Figs. 2A and B), training the model with noisy calls (red lines) significantly improved performance when the model was tested with noisy calls (Figs. 2A and B). Interestingly, the models trained on noisy calls sometimes performed slightly worse to categorize clean calls compared to the models that were trained on clean calls (e.g., *Twitter* and *Chut*, see data point marked ‘∞’ in Figs. 2A, B).

**Figure 2:**
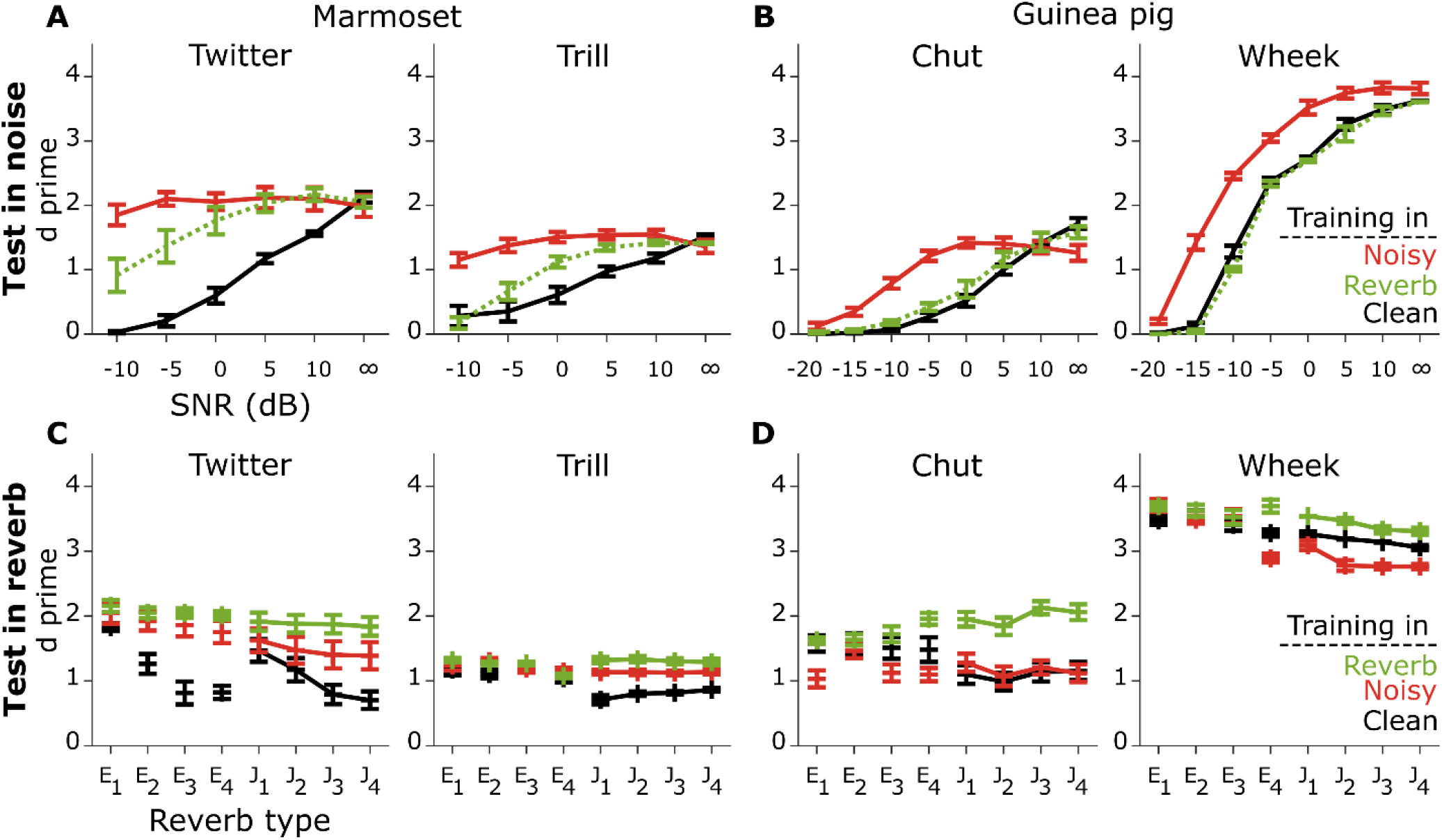
Condition-specific training improved performance but benefits were typically limited to the same condition. (**A, B**) Model performance was quantified by the sensitivity metric, d’, in different levels of noise for several types of marmoset calls (**A**) and guinea pig calls (**B**). (**C, D**) Model performance for marmoset (**C**) and guinea pig (**D**) calls in reverberant conditions. Line color corresponds to training condition: black – training with clean calls, red – training with noisy calls, green – training with reverberant calls. Top row corresponds to testing in noisy conditions and bottom row to testing in reverberant conditions. The benefit was largest when the testing condition matched the training condition (see Table 1 for statistics). Lines correspond to means and error bars denote ±1 s.e.m. Statistics are reported in Table 1.

**Table 1:** Effect of training relative to the baseline model.

| Mechanism<br>Testing | Noisy training |  | Reverberation training |  |
| --- | --- | --- | --- | --- |
|  | Marmoset | Guinea pig | Marmoset | Guinea pig |
| Noise | $\chi^2(3) = 145.0$<br>$p < 2.2 \times 10^{-16}$ | $\chi^2(4) = 64.5$<br>$p = 3.2 \times 10^{-13}$ | $\chi^2(3) = 85.7$<br>$p < 2.2 \times 10^{-16}$ | $\chi^2(4) = 5.6$<br>$p = 0.23$ |
| Reverberation | $\chi^2(3) = 81.1$<br>$p < 2.2 \times 10^{-16}$ | $\chi^2(4) = 28.8$<br>$p = 8.4 \times 10^{-6}$ | $\chi^2(3) = 204.4$<br>$p < 2.2 \times 10^{-16}$ | $\chi^2(4) = 71.0$<br>$p = 1.4 \times 10^{-14}$ |

To test whether these benefits translated across conditions, we tested the noise-trained model on calls degraded by reverberation (red lines in Figs. 2C, D) and vice versa (green lines in Figs. 2A, B). Compared to same-condition training, across-condition training led to significantly lower improvement (indicated by lower *χ*^2^ values in Table 1) over the clean-trained model for call types of both species (i.e., marmosets and guinea pigs). In fact, across-condition training sometimes resulted in poorer performance than the original model [e.g., noise-trained *Wheek* model when tested in reverberation, *χ*^2^(1) = 33.9, *p* = 5.9 × 10^−9^]. Overall, these results show that condition-specific training can improve auditory categorization performance, but these benefits do not generalize well to other conditions.

Next, we tested whether FD properties were systematically different for models trained using noisy calls or reverberant calls compared to the model trained using clean calls. We considered the duration, center frequency, bandwidth, threshold, and reduced kurtosis of FDs. We found little systematic differences for either marmoset call types (Fig. S1, Table S1) or guinea pig call types (Fig. S2, Table S2), as indicated by 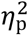 values, which were less than or equal to 0.04. The exceptions were feature duration, which was significantly longer, and threshold, which was significantly lower, for FDs in noise-trained models. Intuitively, an increase in feature duration is expected because integration over longer durations might be necessary to overcome degradation of features by noise. But a reduction (large and significant for marmoset calls, small and significant for guinea pig calls) in FD threshold was not expected because noise is traditionally assumed to increase the mean and reduce the standard deviation of neural activity (Rabinowitz et al., 2011). Overall, the spectrotemporal properties of the FDs were largely unaffected by training in noise or reverberation.

Training in different listening conditions is biologically realistic. In both humans and animals, the acquisition of vocalization categories early in development could conceivably occur in a range of different listening environments including clean, noisy, and reverberant conditions. But our results also illustrate two underlying problems with this approach: 1) features learned during early exposure to a limited set of conditions would not be sufficient for all possible conditions that an individual might encounter over their lifetimes, and 2) a different set of features necessary for each listening condition would combinatorially increase the number of features that need to be represented in cortex. An alternative approach would be to learn a small set of features in clean or a limited set of conditions and use other adaptive neural mechanisms to either modify noisy inputs to simulate clean inputs or modify feature properties to handle noisy inputs. As a first step towards modeling such adaptive mechanisms, we characterized the effects of additive white Gaussian noise on activity at the spectrotemporal input layer, which serves as the input to the FD layer. We also characterized the effects of noise on the *V*_*m*_ responses of FDs, which were trained in clean conditions.

### Noise differentially affected amplitude distributions at feature-detector input and response

The distributions of response amplitudes across all frequency and time bins at the spectrotemporal layer, i.e., the FD inputs (Fig. 3B), and across all time bins of the FD *V*_*m*_ responses (Fig. 3C) are shown in Figure 3. Similar to the effects of noise addition to the acoustic waveform (i.e., increase in mean and decrease in standard deviation) (Rabinowitz et al., 2011), noise increased the mean [2.6 (clean), 3.0 (0 dB), and 3.5 (−10 dB)] and reduced the standard deviation [2. 7 (clean), 2.0 (0 dB), and 1.9 (−10 dB)] of the dense spectrotemporal layer (FD inputs, red and green lines in Fig. 3B). The FD *V*_*m*_ response also showed an increase in the mean amplitude (rightward shift in distributions, Fig. 3C) and a decrease in its standard deviation (smaller spread in distributions, Fig. 3C) with increasing noise level. However, what is relevant for categorization is whether a given feature was detected, i.e., whether the maximum value of the FD *V*_*m*_ response (triangles) exceeded the threshold of that FD. We observed that the maximal FD *V*_*m*_ response in fact decreased with increasing noise level. This is likely because the FD *V*_*m*_ response is the extent to which a given feature ‘matches’ (using the max. cross-correlation value as a metric) the input, and the max. cross-correlation values progressively decreased with increasing noise level. We had earlier commented on our observation that features trained in noisy conditions exhibited lower thresholds than those trained in clean conditions. This decreased threshold likely compensates for the lower maximal values in the FD *V*_*m*_ response distributions, thereby resulting in a supra-threshold FD output.

**Figure 3:**
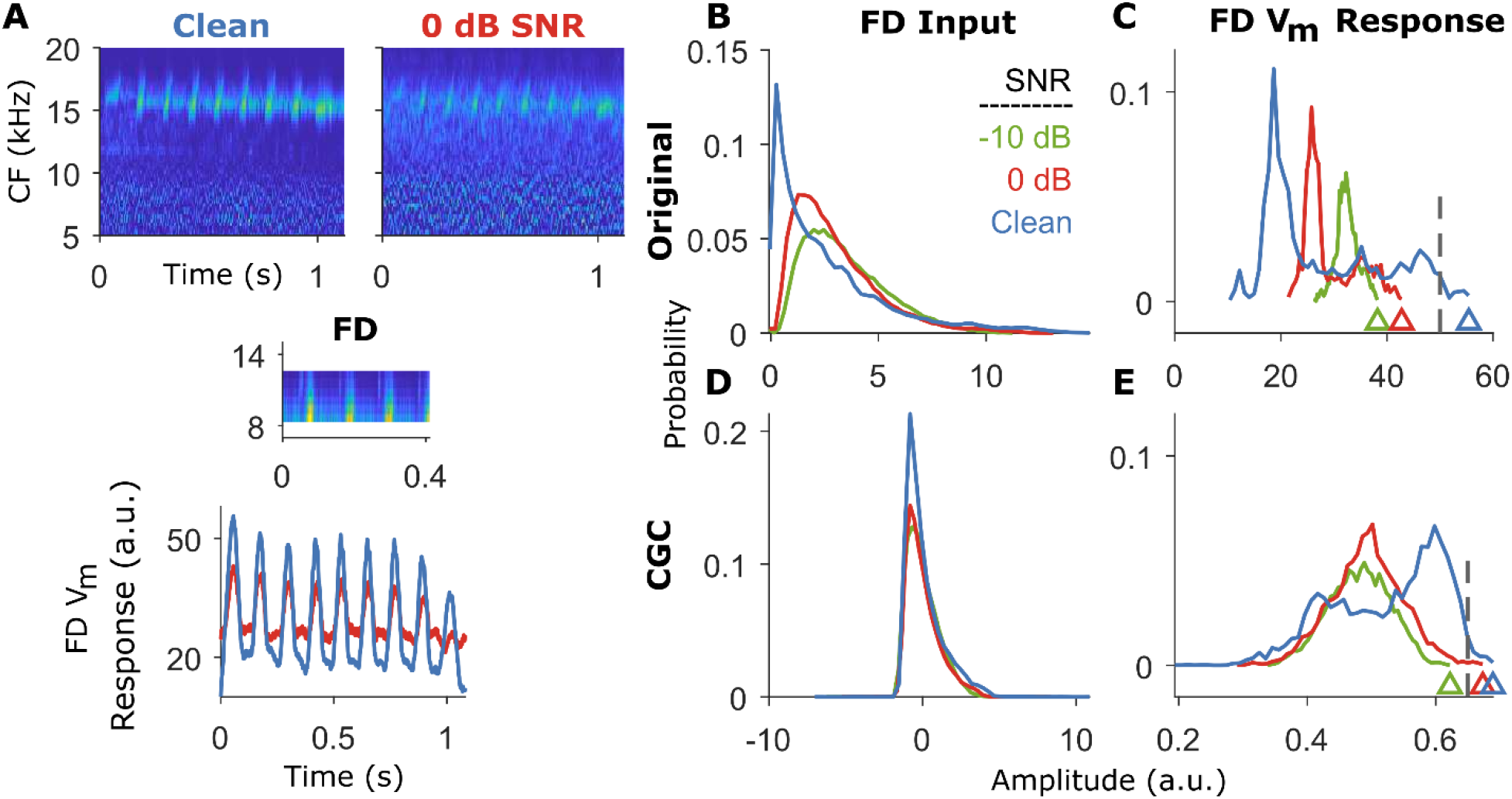
Noise differentially affected amplitude distributions of the input and *V*_*m*_ response of a FD. (**A**) *top*, Example cochleagrams for a marmoset twitter call in clean and 0 dB SNR conditions. *Middle*, example cochleagram of a most informative feature for categorizing twitters. *Bottom, t*he *V*_*m*_ response of an example FD had a lower standard deviation in the noisy condition (red) compared to the clean condition (blue). (**B-C**) Amplitude distribution of the input (**B**) and response (**C**) of the FD in **A** (without contrast gain control) for three different SNRs. Dashed line in **C** represents the threshold of the FD learned during training. Triangles in **C** denote the maximum FD response value, which is the quantity subject to thresholding. (**D-E**) Same as **B-C** but with contrast gain control included in the model. CF, Characteristic frequency; CGC, Contrast gain control; FD, Feature detector.

This observation of obtaining a supra-threshold FD output in noisy conditions by lowering the threshold in noise-trained features led us to explore two alternative strategies to counter these effects of noise to improve model performance in noise (and reverberation) with a single set of FDs trained in the clean condition. First, we implemented contrast gain control, which normalizes the input to FDs such that the mean and standard deviation do not change with noise level (Fig. 3D). Note that this is an upper bound on stimulus-contrast restoration (i.e., complete restoration within the receptive fields of FDs) as neurophysiological data show that the auditory system only partially restores stimulus contrast (Wen et al., 2009; Willmore et al., 2014). Second, because noise decreases the maximum value of the FD *V*_*m*_ response (for both the original model in Fig. 3C and the model with contrast gain control in Fig. 3E), the threshold can be dynamically varied based on the level of noise to improve feature detection in noise. In the following, we test the efficacy of these two adaptive mechanisms.

### Contrast gain control improved model performance in noise and reverberation

In the original model (i.e., the model without contrast gain control), the FD STRF was demeaned and normalized to have unity variance. This STRF was cross-correlated with the input cochleagram, and as such, there was no stimulus-dependent contrast gain adjustment to the input cochleagram in the original model. To computationally implement contrast gain contrast, we demeaned both the input cochleagram (within the FD bandwidth) and the FD STRF, and normalized both to have unity standard deviation. After this normalization, FD input amplitudes were largely overlapping at all tested noise levels (Fig. 3D), and the range of FD *V*_*m*_ responses was limited between -1 and 1. Compared to the model without gain control, the FD *V*_*m*_ response distributions for the gain control model showed increased overlap, and the maximal value of the *V*_*m*_ response distributions crossed the threshold at more adverse noise levels (Fig. 3E). When we tested categorization performance, we saw that contrast gain control improved model performance both in noise and in reverberation for most call types (Fig. 4, Table 2). However, the trends of these benefits were rather heterogenous (e.g., high improvement at low SNRs for *Twitter* but at high SNRs for *Chut*, and little benefit for *Wheek* in either noise or reverberation). Overall, these results show that contrast gain control can improve auditory categorization in noise and in reverberation in a computational model, but that even a perfect implementation of contrast gain control is insufficient to obtain significant benefits across all stimulus types.

**Figure 4:**
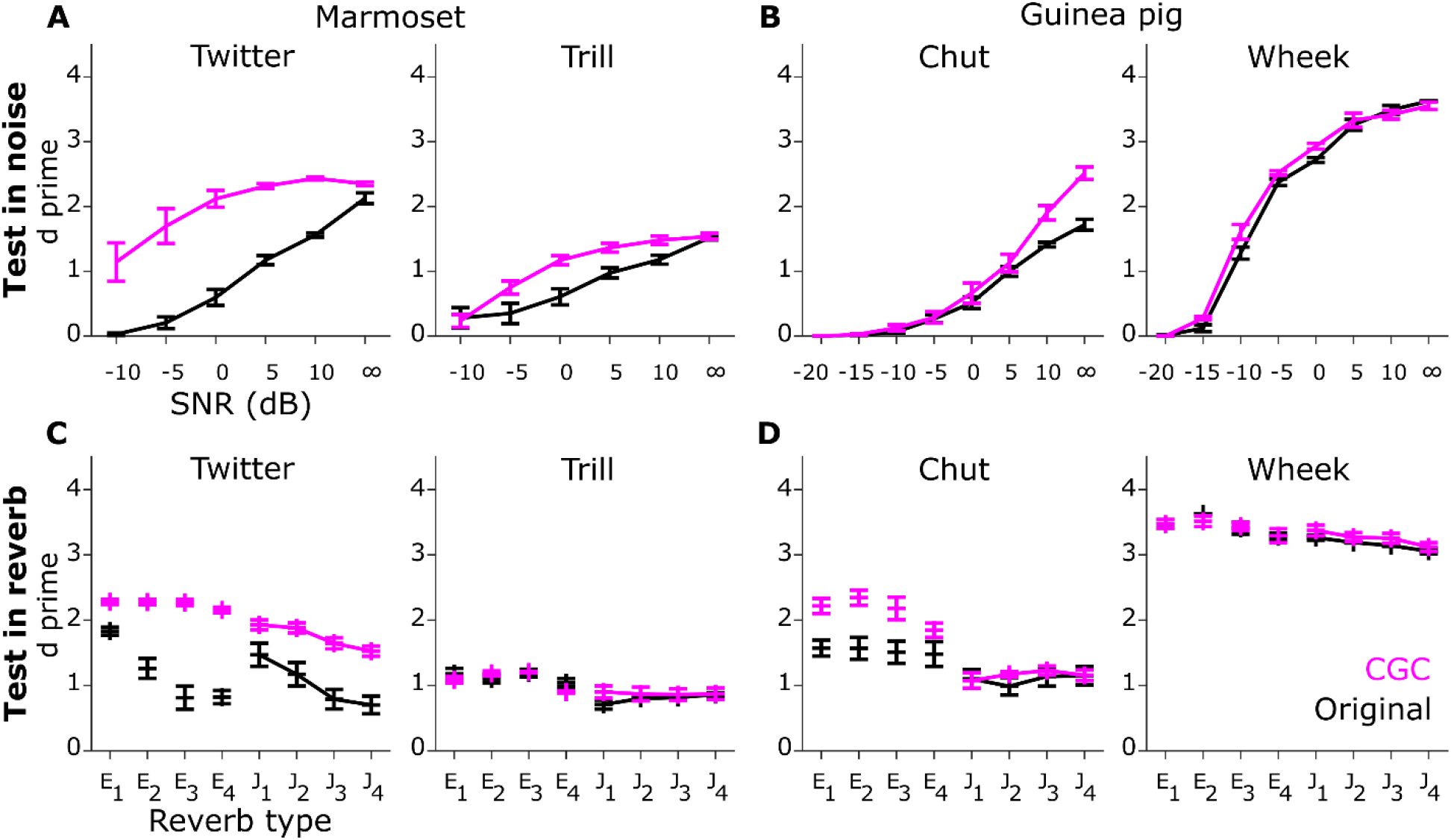
Contrast gain control improved model performance in noise and reverberation. Same format as Fig. 2. Black and magenta lines correspond to the model without and with contrast gain control, respectively. Lines correspond to means and error bars denote ±1 s.e.m. Statistics are reported in Table 2.

**Table 2:**
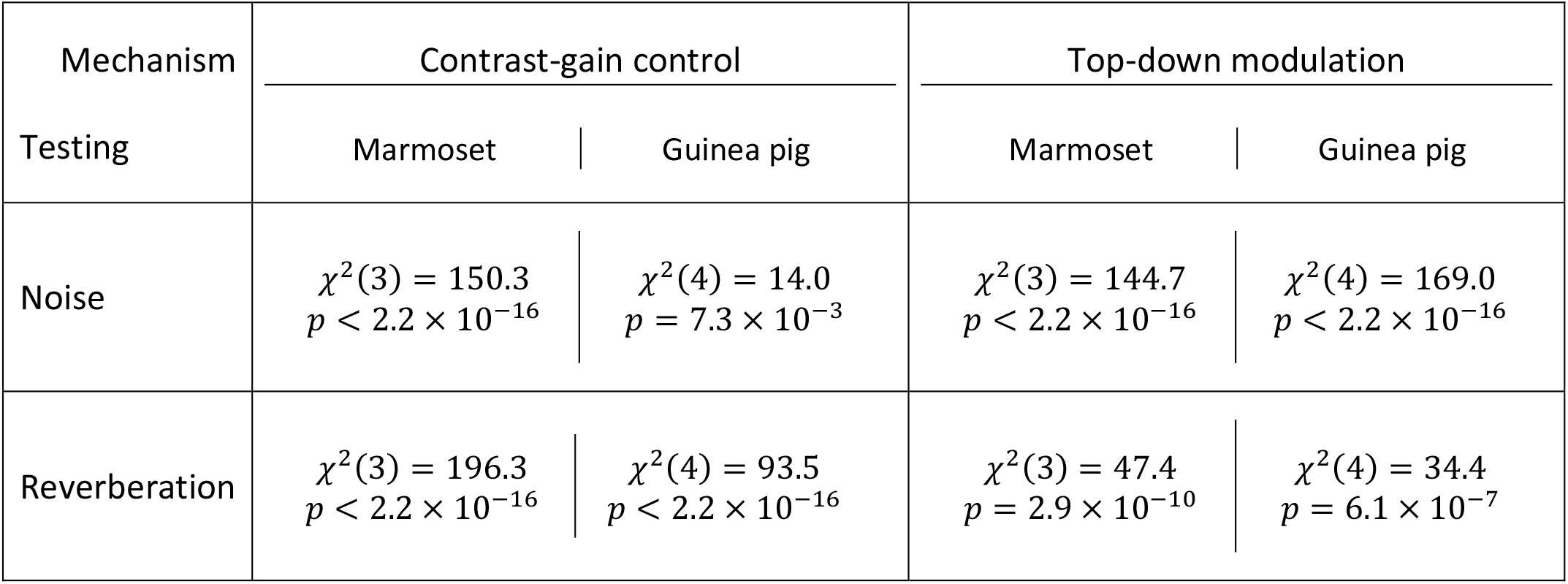
Effect of contrast-gain control and top-down modulation on model performance relative to the baseline model.

### Model performance improved with top-down excitability modulation, but the magnitude of modulation scaled with SNR and not with reverberation strength

Next, we evaluated the merit of top-down influences to improve model performance. As mentioned earlier, noise reduced the maximum *V*_*m*_ response value of a FD. But crucially, this reduction occurred for both within-class as well as outside-class calls for a FD (Fig. 5A). Therefore, even though both within-class and outside-class distributions of maximum FD *V*_*m*_ responses were shifted down to lower values, these distributions retained some degree of separation, and optimal performance could theoretically be obtained by scaling down the FD threshold appropriately. We estimated the optimal threshold ratio for each condition (each SNR or reverberant condition) as the ratio value that maximized classification in that condition (quantified using the mutual information between true and predicted call types). This optimal threshold ratio approximately linearly scaled with SNR for both marmoset and guinea pig calls (Fig. 5B and 5D) and was more or less consistent across call types for each species. This was not the case for different reverberation conditions, however (Fig. 5C and 5E); in this case, the optimal threshold ratio was about the same across many tested reverberant conditions. Biologically, the reduction in threshold with noise level could be accomplished by increasing the excitability of FD neurons, i.e., by reducing the distance between each neuron’s resting membrane potential and spike threshold. In a neural circuit, this could be accomplished by disinhibiting these neurons, perhaps through the activation of specific cortical cell types by cholinergic or noradrenergic inputs (Froemke et al., 2007; Kuchibhotla et al., 2017; Letzkus et al., 2011; Maness et al., 2022; Martins and Froemke, 2015; Pi et al., 2013).

**Figure 5:**
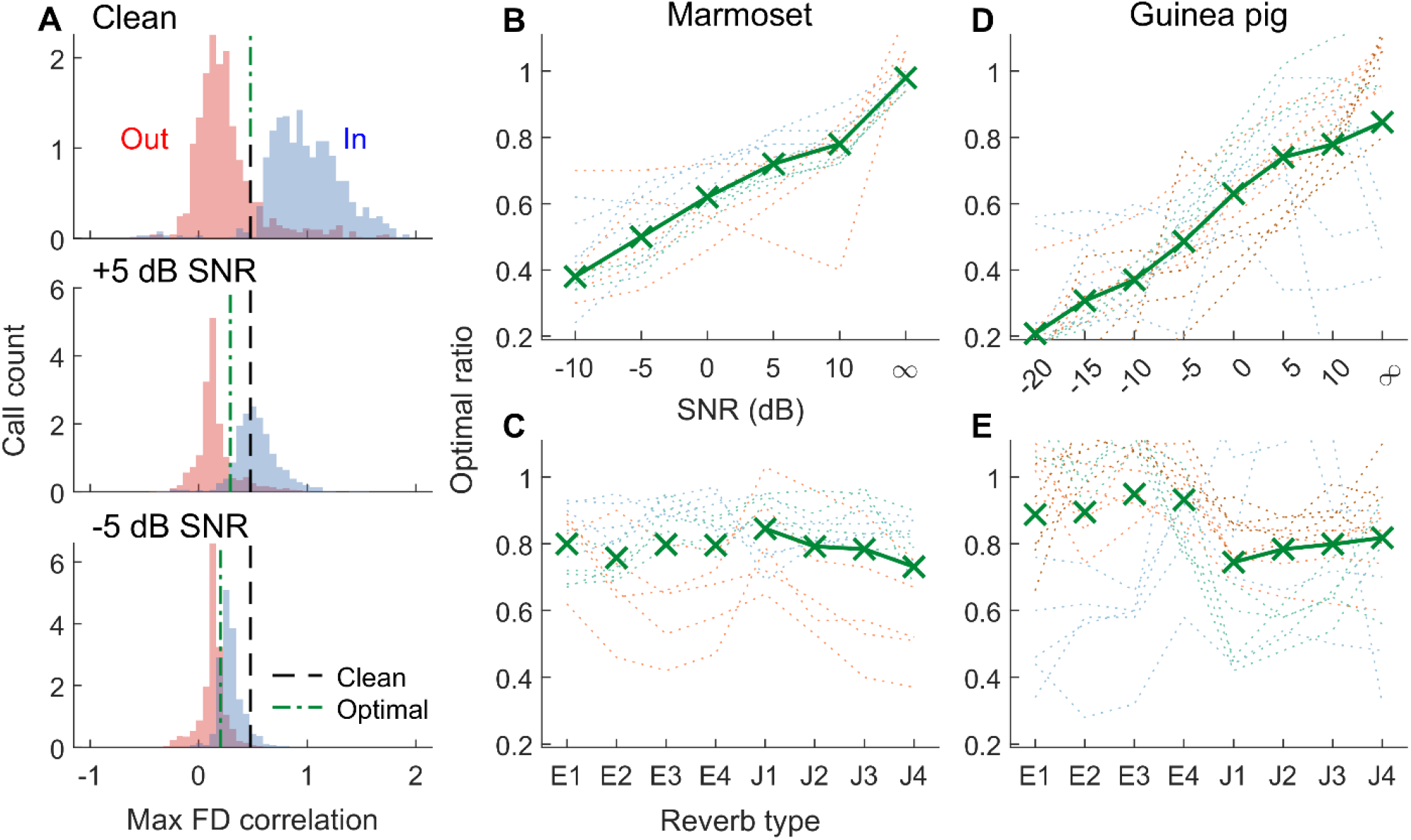
The distribution of feature-detector responses linearly scaled with SNR but not with the strength of reverberation. (**A**) Distributions of *V*_*m*_ response maximum for a *Twitter* FD for 500 within-class (blue) and 500 outside-class (red) calls in clean (top), +5 dB SNR (middle), and -5 dB SNR (bottom) conditions. Black dashed line denotes the threshold learned during clean training; green dash-dotted line denotes the optimal threshold in each SNR condition. **(B, D)** The optimal threshold values for different SNRs for marmoset (**B**) and guinea pig (**D**) call types. (**C, E**) The optimal threshold values for different reverberant conditions for marmoset (**C**) and guinea pig (**E**) call types. In **B-E**, thin dotted lines represent individual call types and solid green lines with symbols denote the median value across all call types.

Next, we quantified model performance as a result of top-down modulation of FDs. Model performance improved significantly and consistently for all call types in the SNR conditions (Fig. 6, Table 2), and typically benefits scaled with the magnitude of top-down modulation (i.e., optimal ratio in Fig. 5B). Model performance improved but to a lesser extent for reverberant conditions (lower *χ*^*2*^ values in Table 2. In summary, these results show that top-down modulation of FDs improves model performance, but these benefits are greater in noisy conditions than in reverberant conditions.

**Figure 6:**
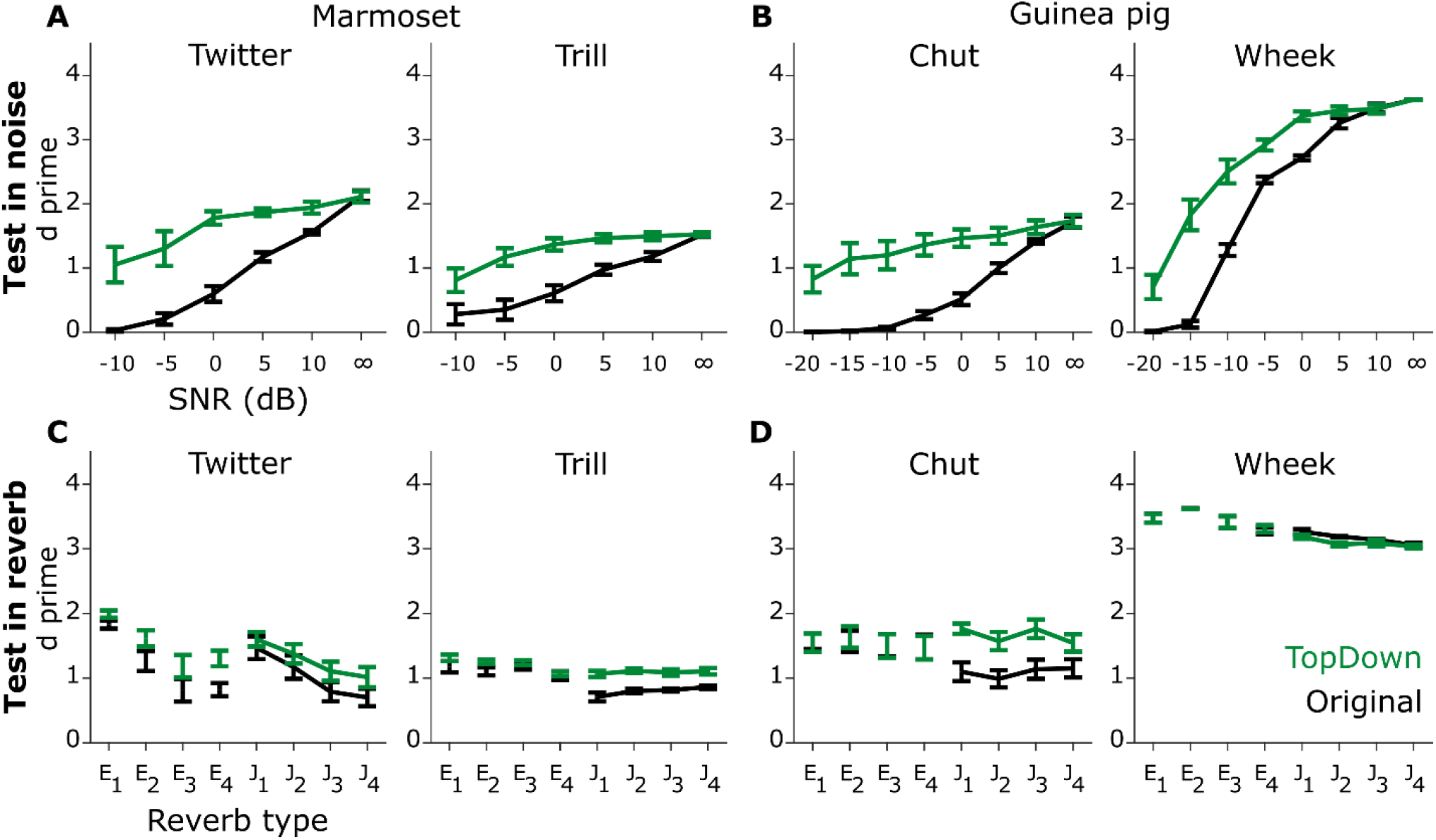
Top-down excitability control significantly improved model performance in noise and less so in reverberation. Same format as Fig. 2. Black and green lines correspond to the model without and with top-down modulation of FD excitability respectively. Lines correspond to means and error bars denote ±1 s.e.m. Statistics are reported in Table 2. Optimal threshold ratios in Fig. 5B were used to implement top-down modulation across conditions.

### Biologically feasible implementation of contrast gain control and top-down modulation of excitability

The adaptive model implementations presented so far are engineering solutions – for example, we implemented perfect contrast gain control by measuring the response distributions of the spectrotemporal layer and normalizing the responses using the mean and standard deviation. However, such implementations may not be biologically realistic. For example, contrast gain control in the auditory system partially restores stimulus contrast at a neuronal level over multiple processing stages, whereas our contrast gain control implementation aimed to restore contrast completely for each FD at a single stage. To determine the extent to which biologically feasible implementations of the mechanisms presented above aid model performance in noise and reverberation, we explored the following alternate implementations (Fig. 7). For contrast gain control, instead of modifying cochleagrams (zero mean and unity variance) in a boxcar manner (matched to the bandwidth and duration of the FD) for each time step of the cross-correlation operation, we subtracted the mean of the entire cochleagram and normalized it to have unity variance. This way, contrast restoration was not optimized to within the bandwidth and duration of individual FDs. Such an operation could be performed in neural circuits by widely tuned inhibitory neurons that have been implicated in contrast gain control in the visual system (Atallah et al., 2012; Markram et al., 2004), although the auditory system may have different implementations (Cooke et al., 2020). We retrained the model with this cochleagram normalization to learn FDs. Similar to the previous model with contrast gain control, this modified implementation of contrast gain control also led to significantly better performance than the original model (i.e., without any contrast gain control, Fig. 7C).

**Figure 7:**
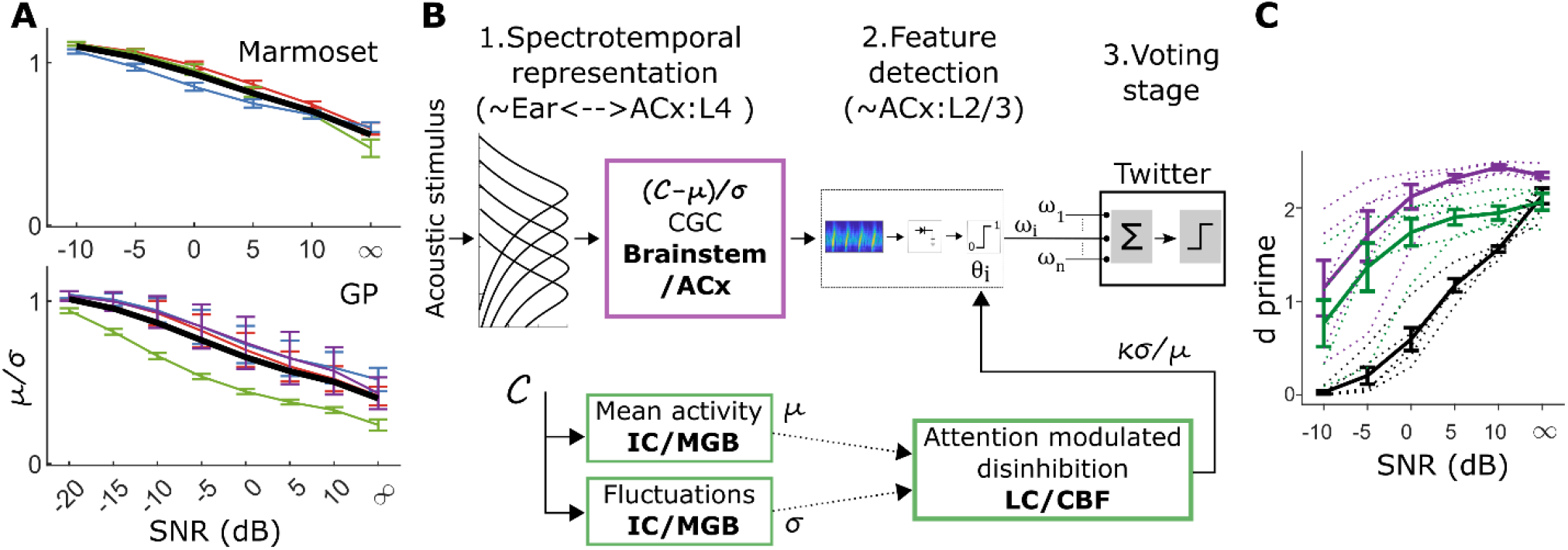
Biologically feasible implementation of contrast gain and top-down control for noisy conditions. **(A)** The ratio of the mean (μ) to standard deviation (σ) of the overall cochleagram at different SNRs for marmoset (top) and guinea pig (bottom) call types. Colored thin lines represent the mean (± s.e.m.) for μ/σ for individual call types; thick black lines indicate the ***median*. (B)** *Top*, contrast gain control can be implemented by demeaning the cochleagram and normalizing it by its standard deviation. *Bottom*, top-down control can be implemented by scaling the feature-detector threshold by a factor proportional to the ratio of standard deviation to mean of the cochleagram. **(C)** Model performance to classify *Twitter* calls from other call types was significantly better using biologically feasible contrast-gain control [*χ*^2^(2) = 70.1, *p* = 5.8 × 10^−16^] as well as top-down modulation [*χ*^2^(2) = 36.6, *p* = 1.4 × 10^−9^]compared to the original model.

To implement top-down control, we had previously used knowledge of stimulus SNR, which is available to the experimenter but is not accessible to biological neural circuits. Therefore, we first asked whether the mean (μ) and standard deviation (σ) of cochleagrams can be used to estimate the acoustic SNR. The mean can be estimated from the population activity of neurons at multiple stages in the auditory pathway. The standard deviation (or modulation) can be estimated biologically by neurons in the inferior colliculus, whose responses have been shown to be sensitive to temporal modulations (Joris et al., 2004; Krishna and Semple, 2000; Nelson and Carney, 2007). We found the ratio of μ over σ to be (negatively) correlated with the acoustic SNR (Fig. 7A). Next, we learned the optimal threshold ratio (κ) based on the μ/σ of cochleagrams, thus doing away with the need to use the acoustic SNR. A single κ (median across all calls) was estimated for all call types of each species (marmoset or guinea pig). To demonstrate feasibility, for a test call type (marmoset *Twitter* call), we used κ to scale the threshold of FDs for individual cochleagrams. Similar to the top-down model, the performance of this biologically feasible model was significantly better than the original model (Fig. 7C).

### Comparison of model performance with behavior

Finally, we compared the performance of the proposed models with guinea pig behavioral performance (Fig. 8). Guinea pigs were trained on a go/no-go task using only clean calls, where the target/distractor call pair was either chut/rumble or wheek/whine. Guinea pig call categorization was then tested in noisy conditions at different SNRs. Details of the behavioral paradigm have been described previously (Kar et al., 2022). Note that guinea pig behavior is governed by two underlying processes – the recognition of the correct call category and the expression of that recognition by performing an operant action. The model only captures the former process, and while other factors such as motivation and attention might influence guinea pigs’ operant behavior, the model perfectly reports the recognized call category. Thus, the models are expected to over-perform when compared to guinea pig behavior. The performance of the original model without adaptive mechanisms matched (for *wheeks*) or was worse (for *chuts*) than guinea pig behavioral performance in noisy conditions (Fig. 8). But all extended models performed better than guinea pigs for categorizing *wheek* calls from distractors. For categorizing *chut* calls, the model with top-down modulation matched behavior at negative SNRs whereas the model with contrast gain control matched behavior at the two most favorable SNRs. These heterogeneous trends suggest that the contributions of these different mechanisms may depend on the call spectrotemporal properties (tonal calls such the *wheek* vs. short-duration noise-like calls such as the *chut*) as well as their interaction with noise level. Taken together, these results suggest that a combination of adaptive mechanisms is necessary for the hierarchical model to recapitulate guinea pig behavioral performance for call categorization in noise.

**Figure 8:**
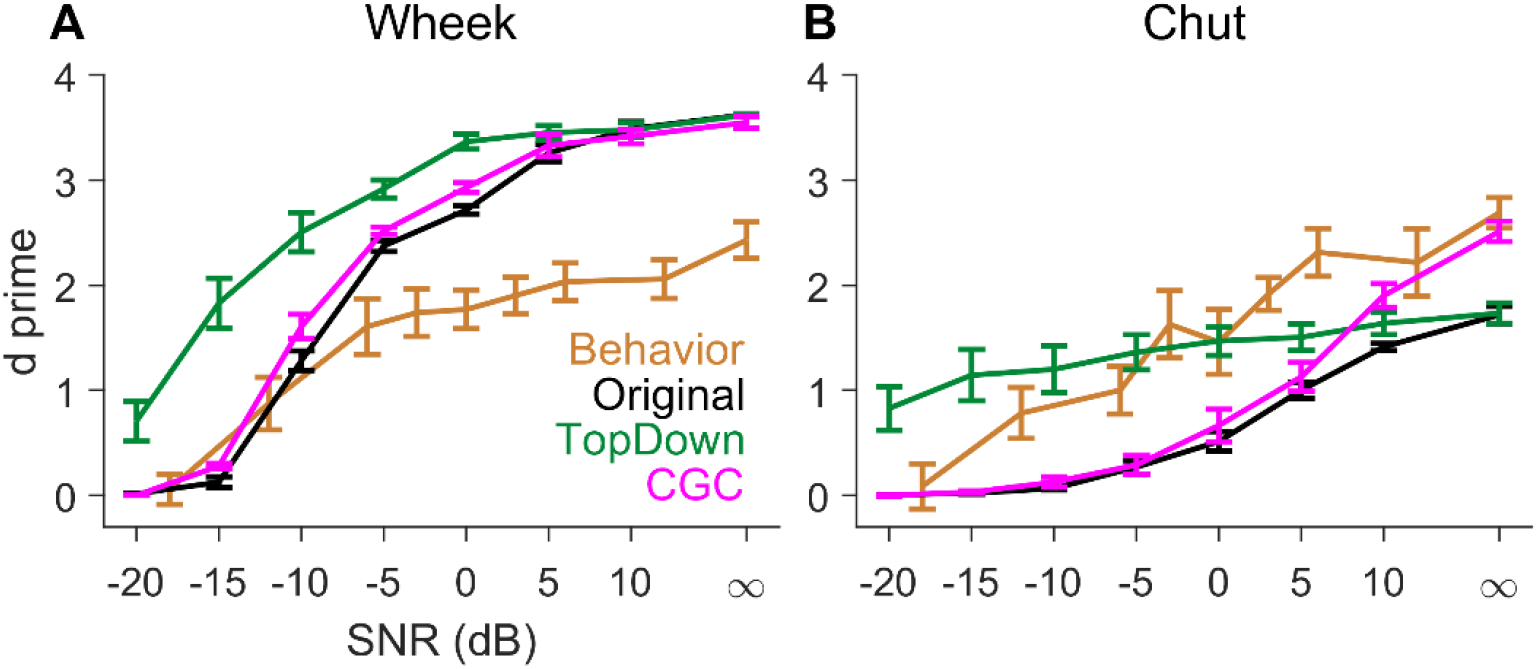
Comparing the performance of various models with the behavioral performance of guinea pigs. **(A-B)** Performance of various models and guinea pigs (behavior) for wheeks (**A**) and chuts (**B**). Guinea pigs were trained on a go/no-go task, where the target/distractor pair was either wheek (go)/whine or chut (go)/purr. The performance metric for guinea pig behavior was determined as d-prime = *norminv*(hit rate) – *norminv*(false alarm), where *norminv* is the inverse Normal cumulative distribution function.

## Discussion

In this study, we explored biologically realistic extensions to a hierarchical model for auditory categorization to improve its performance in realistic listening conditions. We tested three mechanisms including task-specific training, contrast-gain control, and top-down modulation of FD sensitivity. All three mechanisms improved model performance when tested in noisy and reverberant listening environments; however, the trends and magnitude of improvement varied across mechanisms. We demonstrated that the proposed contrast-gain control and top-down modulation mechanisms can be implemented by biological circuits. Finally, comparison of model performance with guinea pig behavioral performance revealed that these mechanisms could be employed in a flexible manner to accomplish call categorization based on both task demands and call spectrotemporal properties.

### Task-specific training led to little systematic differences in the spectrotemporal properties of feature detectors

Consistent with previous studies, our results showed that task-specific training can improve auditory categorization performance in noisy and in reverberant conditions. The benefit was typically the greater at more adverse conditions (e.g., between -10- and 0-dB SNR, and for J1-J4, which were the strongest reverberant conditions). Interestingly, the noise-trained model frequently showed non-monotonic performance curves as a function of SNR and performed worse in clean conditions than models trained on clean calls (e.g., *Twitter, Chut*). While small amount of noise during training can have a regularization effect to improve generalizability of models (Bishop, 1995), uniformly using all SNRs during training likely led to overfitting of the model to noisy conditions. Overfitting may be avoided by appropriately weighing SNRs during training by following SNRs that animals and humans naturally experience.

Even though task-specific training improved model performance in the same task, the benefits were significantly reduced or sometimes negative when training and testing conditions were different. Furthermore, task-specific training did not lead to any systematic changes in spectrotemporal properties of FDs (except for duration for noise-trained models for marmoset calls). Therefore, task-specific training is an inefficient way to categorize sounds.

### Contrast-gain control improved model performance, but benefits varied across call types

Contrast gain control is ubiquitous in both visual and auditory pathways, and more specifically, it has been reported along the auditory pathway from the auditory nerve (Wen et al., 2009) to the auditory cortex (Lohse et al., 2020). Contrast gain control can tune the dynamic range of individual neurons to the statistics of the incoming sound, which can aid in adverse listening conditions. For example, additive noise increases the mean power of the signal while reducing its variance (Willmore et al., 2014). The effect of reverberation on communication signals such as speech and vocalization is also similar as reverberation smears the temporal envelope thus reducing the contrast in the signal (Wang and Brown, 2006). Contrast gain control significantly improved model performance both in noise and in reverberation. However, the trends of improvement were quite heterogeneous across call types, suggesting that the benefits of contrast gain control depend on the spectrotemporal properties of the signal. Future studies are needed to understand what features determine the benefit due to contrast gain control.

### Top-down modulation led to divergent benefits in noise and in reverberation

Attention plays a critical role in shaping perception in challenging environments across modalities (Shinn-Cunningham, 2008). Effects of attention are thought to be mediated by top-down modulation, which can shape the encoding of afferent input along the auditory pathway (Fu et al., 2014; Kerlin et al., 2010). One such pathway involves the vasoactive intestinal polypeptide (VIP)-expressing interneurons that specialize in disinhibition of excitatory cortical principal cells by inhibiting somatostatin and parvalbumin-expressing interneurons (which inhibit principal cells) (Pi et al., 2013). Since VIP-expressing interneurons are activated by cholinergic inputs (at least in the primary visual cortex), which correlate with fluctuation in pupil size (Reimer et al., 2016), this pathway likely captures aspects of effortful listening, which is also known to correlate with pupil size (Peelle, 2018; Zekveld et al., 2010).

We modeled such top-down feedback control by scaling the excitability of FDs to optimally improve task performance in noise and in reverberation. Interestingly, the optimal scale was proportional to noise level, but it was nearly constant across different reverberant conditions. Similarly, benefits of top-down control were greater in noisy conditions than in reverberant conditions (as indicated by *χ*^*2*^ values in Table 2). One interpretation of this result is that increased listening effort is not beneficial in reverberant environments. While unexpected, this interpretation is supported by psychoacoustic studies in humans that demonstrate that listening effort scales with noise level but not with the strength of reverberation (McCloy et al., 2017; Picou et al., 2016; Prodi and Visentin, 2022). These results suggest that top-down attentional mechanisms are primarily beneficial in noisy environments, but these benefits are significantly reduced in reverberant environments.

### Biological feasibility and underlying neural circuits

While contrast gain control has been documented along several regions of the auditory pathways (Angeloni et al., 2021; Barbour and Wang, 2003; Dean et al., 2005; Lohse et al., 2020; Rabinowitz et al., 2011; Watkins and Barbour, 2008; Wen et al., 2009), the underlying neural circuits are not settled. Unlike the visual system, neither shunting inhibition by parvalbumin-positive interneurons nor fluctuation in membrane potential seem to contribute to contrast gain control in the mice auditory cortex (Cooke et al., 2020). Another candidate mechanism that may contribute to contrast gain control is synaptic depression, by which neurons adapt to steady backgrounds (e.g., in noisy or reverberant conditions), thus maintaining their dynamic range to respond to transient signals (e.g., onset of the next syllable in ongoing speech) (David and Shamma, 2013; Mesgarani et al., 2014; Trussell, 1999). Explicitly incorporating synaptic depression would improve the biologic realizability of the computational model.

Several pathways have been shown to modulate the excitability of cortical neurons, which can improve the neural coding of communication sounds, thereby leading to benefits similar to that seen in our model. For example, vasoactive-intestinal-peptide-expressing neurons, which are recruited by reinforcement signals (e.g., reward and punishment during learning), suppress subtypes of inhibitory interneurons, thus, in turn, disinhibiting principal neurons in the cortex during task engagement (Pi et al., 2013). Two other relevant pathways that also modulate cortical excitability involve the noradrenergic locus coeruleus (Maness et al., 2022; Martins and Froemke, 2015) and the cholinergic basal forebrain (Froemke et al., 2007; Kuchibhotla et al., 2017; Letzkus et al., 2011; Pi et al., 2013). While both these pathways generally modulate auditory cortical activity globally (i.e., in a frequency-independent manner), which is similar to our implementation of top-down modulation (e.g., FDs for all calls were similarly modulated at the same noise level or reverberation condition), it remains to be seen whether other top-down mechanisms exist that are optimized to the spectrotemporal properties of the incoming sound.

In conclusion, we have extended a versatile auditory-categorization model, whose strengths include its straightforward biological interpretability and efficient trainability, to include adaptive neural mechanisms, and showed that mechanisms such as contrast-gain control and top-down feedback control can improve auditory categorization performance in challenging listening environments.

## Materials and Methods

### Stimuli

All procedures followed NIH-issued guidelines for the care and use of laboratory animals. The model was trained to perform an auditory categorization task in which it discriminated one call type from other conspecific call types. Two sets of vocalization stimuli were used, including calls from marmosets and guinea pigs, two highly vocal and social species. Marmoset calls have been described in detail in a previous study (Agamaite et al., 2015) and have been used in the previous version of the model (Liu et al., 2019). Briefly, these calls were recorded from eight adult marmosets of either sex living in a marmoset colony using an array of directional microphones. Guinea pig calls were primarily from four male and one female guinea pigs. Male guinea pigs were placed in pairs in a sound-attenuating booth, sometimes in two different chambers separated by an acrylic divider (Montes-Lourido et al., 2022). A directional condenser microphone, suspended from the sound booth ceiling, was used to record these vocalizations. To record *wheek* calls, the microphone was placed outside the guinea pig cages in the colony using a tripod. Guinea pig calls were recorded using Sound Analysis Pro 2011, sampled at 48 kHz, and manually curated using Praat (Boersma, 2001).

Noisy calls were generated by adding white Gaussian noise to calls. Reverberant calls were generated using eight different impulse responses. The strength of reverberation is typically quantified by the T30 metric, which indicates the time it takes for signal energy to decay to 30 dB below the original value. Four of these impulse responses (denoted by J1-J4), have been previously used for human speech perception studies (Jørgensen and Dau, 2011). These impulse responses were originally generated using Odean (Christensen, 1999) and had the following T30s – 128 (J1), 236 (J2), 461 (J3), and 644 (J4) ms. The other four impulse responses were impulse responses corresponded to the following naturalistic environments – snow site (E1, T30 = 7 ms), plastic bin (E2, 57 ms), living room (E3, 81 ms), and forest (E4, 124 ms).

### Model architecture

The current model is similar to the model we have previously used, but there are several differences. Briefly, the model is hierarchical and has three stages. The first stage is a spectrotemporal stage that uses a phenomenological model (Zilany et al., 2014) of the auditory periphery to generate the cochleagram, which is a physiologically accurate spectrotemporal representation of the stimulus. Inner hair cell voltage was used as the response instead of the auditory nerve fiber firing rate (which was used in the previous model) to achieve faster implementation. Cochleagrams were generated at a sampling frequency of 1 kHz and the center frequencies spanned 200 Hz to 20 kHz in 0.1-octave steps. Marmoset calls were processed using a marmoset head-related transfer function (Slee and Young, 2010), which was estimated using the GRABIT code (Doke, 2022). Cochlear tuning in the periphery model was set to “*human*” for marmoset cochleagrams and to “*cat*” for guinea pig cochleagrams. The overall stimulus level was of clean and degraded calls was set to 65 dB SPL. In the second stage, the cochleagram is filtered by the STRF of a feature detector (FD) to obtain its membrane potential response or *V*_*m*_ response, following which a threshold is applied to the maximum of the filter output to obtain a binary output (1 if maximum *V*_*m*_ response > threshold). Finally, outputs of a set of maximally informative FDs are weighted and combined in a voting stage to obtain the final response of the model for a call. The set of maximally informative FDs as well as their thresholds and weights are learned during training, as described next.

The model was trained to classify one call type (within-class) from all other conspecific call types (outside-class). Marmoset models were trained using 500 within-class calls and 500 outside-class calls and tested with a non-overlapping set of within- and outside-class calls (500 calls each). Guinea pig models used 70% of all available calls for training and the remaining for testing. Therefore, the number of calls for training (and testing) were different for different call types (number of training calls for chut=248, rumble=176, wheek=230, whine=300). During training, initial features were generated by randomly segmenting rectangular blocks (i.e., random center frequency, bandwidth, onset time, and duration) of within-class cochleagrams. The number of initial features was 6000 for marmoset call types and 4000 for guinea pig call types.

For each FD, its spectrotemporal pattern was used as the STRF to filter a cochleagram, and the maximum value of the *V*_*m*_ response was used to construct distributions for within-class and outside-class calls. The threshold of the FD was set to the correlation value that maximized the mutual information between the binary output (after applying threshold on the FD *V*_*m*_ response) and the stimulus category (within-class and outside-class). The weight of the FD was set to the log-likelihood ratio of this classification. After estimating the threshold and weight of individual FDs, a greedy search was implemented to estimate a set of maximally informative and least redundant FDs (MIF set) for each model. This was an iterative process, where we sequentially added FDs to the MIF set to increase hit rate without increasing false alarm rate. The maximum number of FDs in an MIF set was set to 20. We trained five instantiations of the model for each call type (by using non-overlapping feature detectors) to assess the reliability of model performance statistically. Model performance was quantified using receiver-operating curve (ROC) analysis; we first estimated the area under the ROC curve (*AUC*) using model output (i.e., output of the voting stage) for within-class and outside-class calls and then estimated the sensitivity index (d’) from this area as: 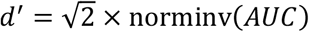, where norminv is the inverse of the Normal cumulative distribution function. The same set of within-class and outside-class calls were used to quantify performance in clean, noisy, and reverberant conditions.

### Feature detector properties

We characterized FDs by estimating several properties such as threshold, center frequency, bandwidth, and duration. In addition, we also estimated the complexity (tailedness) of FDs using reduced kurtosis (i.e., 3 subtracted from the kurtosis, where kurtosis is the ratio of the fourth central moment to the fourth power of the standard deviation). Higher (lower) values for reduced kurtosis indicate a heavy-tailed (light-tailed) distribution relative to the Normal distribution.

### Statistical analysis

All statistical analyses were performed in R (Team, 2013) (version 4.2.1). The effects of various mechanisms were evaluated by constructing linear mixed-effect models (*lme4* package (Bates et al., 2015, p. 4)) and comparing model fits using *anova* (*stats* package). To evaluate the effect of a *mechanism* (four mechanisms in total: noisy training, reverberation training, contrast-gain control, or top-down modulation) on the performance in a testing condition (i.e., in noise or in reverberation), two models were constructed (separately for each species):

1. full model: *dprime ∼ testing_parameter + call_type*mechanism + (1*|*model_id)*

2. null model: *dprime ∼ testing_parameter + call_type + (1*|*model_id)*

where *testing_parameter* was either SNR in dB (interval scale) or reverberation type (nominal scale), *call_type* (nominal scale) was the conspecific call types used for each species, and *model_ID* (a random effect, nominal scale) was the model instantiation index. The term *call_type* was excluded for statistical analysis in Fig. 7 as the data were for only a single marmoset call type (i.e., twitter).

The effect of training on model FD parameters (e.g., duration, CF) was quantified using the F-test as well as the partial eta squared (Richardson, 2011) (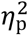, estimated using the *etaSquared* function in the *lsr* package (Navarro, 2015) with type II sum of squares) values, which approximates the fraction of total variance captured by training. A linear model (*lm* in *stats* package) was fit for each combination of species, training condition, and FD property with FD property value as the outcome. Predictors included call type, training condition, and their interaction. Center frequency was log transformed.

## Acknowledgements

This work was supported by funding from the NIH to S. S. (R01DC017141 and R01DC013315). The authors would like to acknowledge Dr. Xiaoqin Wang (Johns Hopkins University) for providing marmoset calls used for modeling. The authors are grateful to Manaswini Kar, Dr. Marianny Pernia, and Kayla Williams for guinea pig behavioral data. This research was supported in part by the University of Pittsburgh Center for Research Computing through the resources provided.

## Author contributions

All authors contributed to the conceptual development of the model. SP implemented all simulations and statistical analyses, building upon a preliminary implementation by STL. SS provided overall guidance, supervision, and funding. SP wrote the manuscript with inputs from SS.

## Supplementary information

**Figure S1.**
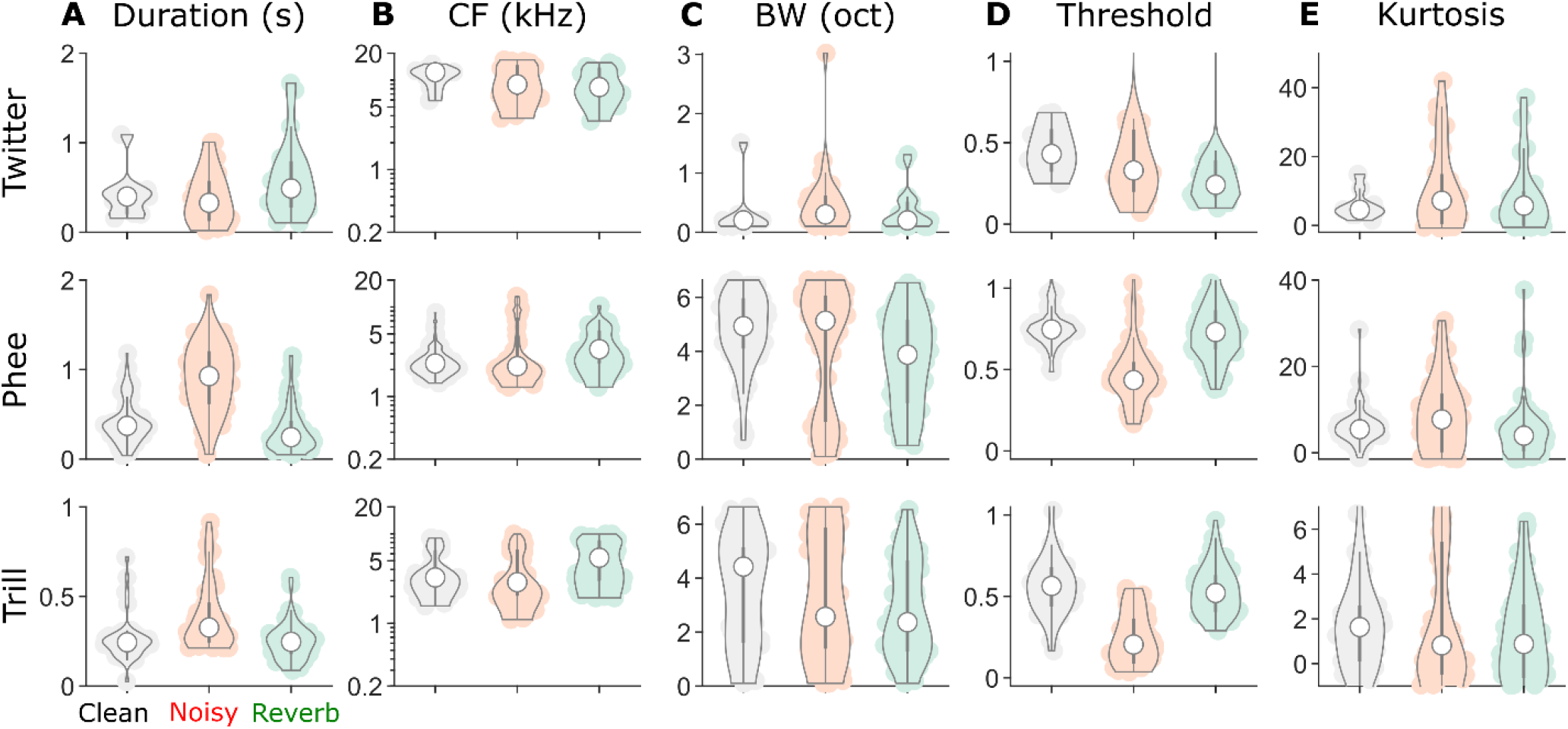
For marmoset call types, properties of FDs were not systematically different across training conditions except for longer duration and lower threshold in noise training. **(A-E)** Distributions of FD properties including duration (**A**), center frequency (**B**), bandwidth (**C**), threshold (**D**), and reduced kurtosis (**E**) of the FD spectrotemporal receptive field for three different marmoset call types (rows). Distributions were constructed using five different model instantiations for each call type. Statistics are reported in Table S1.

**Figure S2.**
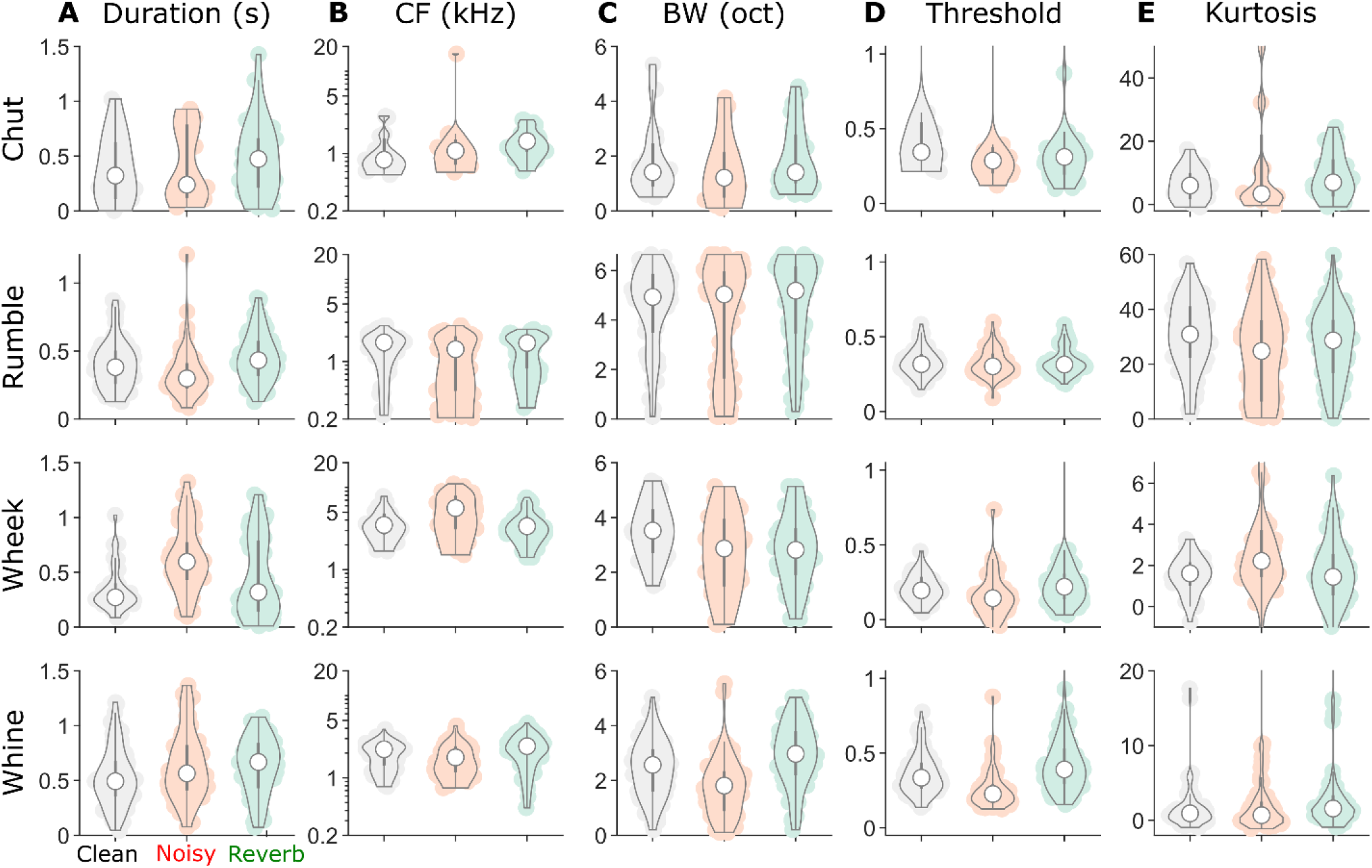
For guinea pig call types, the properties of FDs were not systematically different across training conditions. Same format as Fig S1, but for guinea pig call types. Statistics are reported in Table S2.

**Table S1:** Effect of noisy and reverberation training on FD properties for marmoset calls.

| Training | Duration | CF | Bandwidth | Threshold | Kurtosis |
| --- | --- | --- | --- | --- | --- |
| Noise | $\eta_p^2 = 0.25$<br>$F_{1,629} = 211.4$<br>$p < 2.2 \times 10^{-16}$ | $\eta_p^2 = 0$<br>$F_{1,629} = 1.8$<br>$p = 0.17$ | $\eta_p^2 = 0$<br>$F_{1,629} = 0.01$<br>$p = 0.91$ | $\eta_p^2 = 0.24$<br>$F_{1,629} = 194$<br>$p < 2.2 \times 10^{-16}$ | $\eta_p^2 = 0.03$<br>$F_{1,629} = 16.3$<br>$p = 6 \times 10^{-5}$ |
| Reverb | $\eta_p^2 = 0$<br>$F_{1,710} = 0.4$<br>$p = 0.51$ | $\eta_p^2 = 0.02$<br>$F_{1,710} = 16.9$<br>$p = 4.3 \times 10^{-5}$ | $\eta_p^2 = 0.04$<br>$F_{1,710} = 28.9$<br>$p = 1 \times 10^{-7}$ | $\eta_p^2 = 0$<br>$F_{1,710} = 0$<br>$p < 0.89$ | $\eta_p^2 = 0$<br>$F_{1,710} = 0.1$<br>$p = 0.77$ |

**Table S2:**
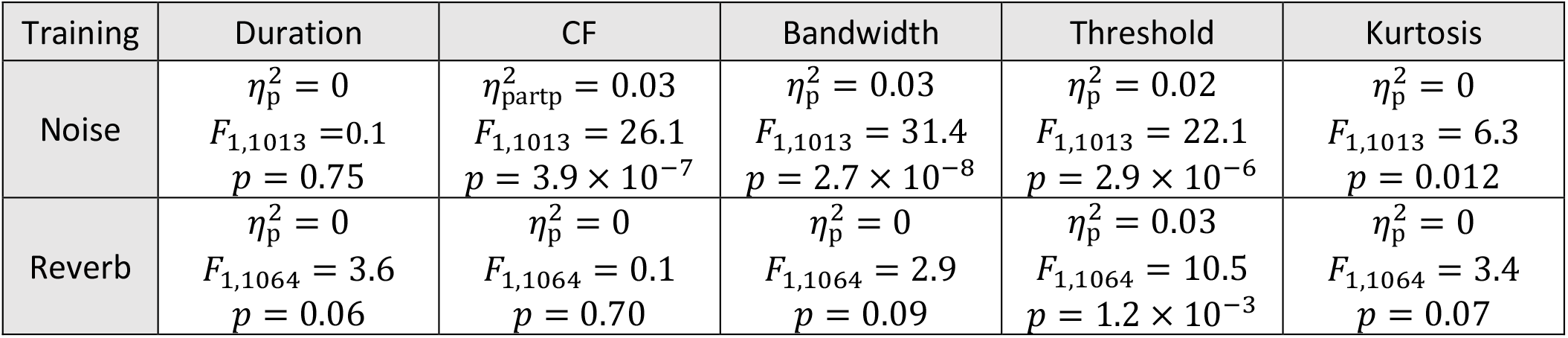
Effect of noisy and reverberation training on FD properties for guinea pig calls.

